# G6PC2 controls glucagon secretion by defining the setpoint for glucose in pancreatic α-cells

**DOI:** 10.1101/2023.05.23.541901

**Authors:** Varun Bahl, Eric Waite, Reut Rifkind, Zenab Hamdan, Catherine Lee May, Elisabetta Manduchi, Benjamin F. Voight, Michelle Y.Y. Lee, Mark Tigue, Nicholas Manuto, the HPAP Consortium, Benjamin Glaser, Dana Avrahami, Klaus H. Kaestner

**Author notes:** Corresponding Authors: Klaus H. Kaestner, Ph.D., M.S., 12-126 Translational Research Center 3400 Civic Center Blvd, Philadelphia, PA 19104-6145, Dana Avrahami, Ph.D., Department of Developmental Biology and Cancer Research, The Hebrew University-Hadassah Medical School, Jerusalem, Israel 91120.

## Abstract

Impaired glucose suppression of glucagon secretion (GSGS) is a hallmark of type 2 diabetes. A critical role for α-cell intrinsic mechanisms in regulating glucagon secretion was previously established through genetic manipulation of the glycolytic enzyme glucokinase (GCK) in mice. Genetic variation at the *G6PC2* locus, encoding an enzyme that opposes GCK, has been reproducibly associated with fasting blood glucose and hemoglobin A1c levels. Here, we find that trait-associated variants in the *G6PC2* promoter are located in open chromatin not just in β− but also in α-cells, and document allele-specific *G6PC2* expression of linked variants in human α– cells. Using α-cell specific gene ablation of *G6pc2* in mice, we show that this gene plays a critical role in controlling glucagon secretion independent of alterations in insulin output, islet hormone content, or islet morphology; findings we confirmed in primary human α-cells. Collectively, our data demonstrate that *G6PC2* impacts glycemic control via its action in α-cells and suggest that *G6PC2* inhibitors could help control blood glucose through a novel, bi-hormonal mechanism.

## Introduction

The endocrine pancreas plays a dominant role in the regulation of glucose homeostasis through the coordinated but opposing actions of glucagon-producing α-cells and insulin-producing β-cells(*1, 2*). Disturbances in this regulatory network result in debilitating diseases such as type 2 diabetes mellitus (T2D), which affects over 425 million people worldwide and is a leading cause of death in many countries(*3*). The “bi-hormonal hypothesis”, published by Unger and Orci almost 50 years ago, proposes that T2D pathophysiology is a combination of insufficient insulin production by β-cells and elevated glucagon release by α-cells(*4*). Consistent with this hypothesis, several studies have illustrated that while patients with T2D exhibit inappropriately low insulin secretion following a meal, plasma glucagon levels also fail to decrease in response to elevated blood glucose, contributing to hyperglycemia(*5–8*).

Given that hyperglucagonemia can exacerbate elevated blood glucose levels, and its correction could provide a therapeutic benefit, there has been a growing interest in elucidating the precise mechanisms governing glucagon release by α-cells. A critical role for α-cell-intrinsic mechanisms has been established through several recent studies in which manipulation of the expression or activity of the glycolytic enzyme glucokinase (GCK) changed the setpoint for glucose-suppression of glucagon secretion (GSGS). For instance, Basco and colleagues showed that mice with α-cell-specific deletion of GCK are unable to suppress glucagon secretion in response to elevated glucose, resulting in hyperglucagonemia(*9*). Another study by Moede and colleagues demonstrated that glucokinase inhibition, activation, or shRNA-mediated gene suppression alters the glucose threshold for glucagon release in single rat α-cells(*10*). In addition, we recently reported that genetic activation of α-cell glucokinase enhances GSGS *in vivo* and *ex vivo* and improves glucagonemia under diabetogenic conditions of high-fat diet feeding(*11*). Altogether, these studies demonstrate that glucose sensing via α-cell glucokinase is a key determinant of glucagon secretion.

The catalytic subunit of the islet-specific glucose-6-phosphatase enzyme (*G6PC2*) opposes the action of glucokinase and creates a futile substrate cycle(*12–14*). Multiple genome wide association studies (GWAS) have linked polymorphisms in *G6PC2* with variations in fasting blood glucose and HbA1c levels with high statistical significance(*15–18*). Molecular studies examining the functional impact of these SNPs on *G6PC2* transcription and splicing have suggested that they directly modulate the expression of *G6PC2* in β-cells(*19*). Animal studies investigating *G6PC2* function *in vivo* and *ex vivo* have demonstrated its role in regulating fasting blood glucose and glucose stimulated insulin secretion (GSIS). Thus, germline *G6pc2* null mice and mice with β-cell-specific ablation of *G6pc2* display significantly reduced fasting blood glucose levels, in support of the notion that *G6PC2* encodes the effector transcript identified by the aforementioned GWAS studies(*12, 13, 20-22*). Moreover, pancreatic islets from *G6pc2* null mice exhibit higher insulin secretion at submaximal glucose concentrations, consistent with a leftward shift of the dose response curve for GSIS(*13*). Collectively, these studies support a role for *G6PC2* in glucose-sensing within β-cells.

Although much effort has been directed toward understanding the consequences of variations in *G6PC2* activity in β-cells, to date no study has analyzed their effect on glucagon secretion in α-cells. When determining cell type specific chromatin accessibility maps of sorted human islet cells(*23*), we noted that glycemic trait-associated SNPs in the *G6PC2* promoter are located in open chromatin regions not just in human β-cells but also in α-cells, suggesting that they might affect both β-cell and α-cell expression of this critical gene. Since glycolytic flux has been shown to affect glucagon secretion(*9–11*), and reduced *G6PC2* levels in β-cells increased insulin secretion(*12, 13*), we hypothesized that reduced *G6PC2* levels would suppress glucagon secretion from α-cells in response to glucose. To test this hypothesis, we derived a new mouse line that permits inducible, α-cell-specific *G6pc2* ablation. Using this model, we provide evidence that G6PC2 in α-cells affects glucagon secretion by modulating the set point for glucose sensing and GSGS. This effect was further demonstrated in an *ex vivo* model of *G6PC2*-deficient human α-cells in pseudo-islets, suggesting a similar role for *G6PC2* in human α-cells and supporting the relevance of *G6PC2* as a therapeutic target to increase insulin secretion while inhibiting glucagon secretion.

## Results

### The islet-specific glucose-6-phosphatase catalytic subunit (*G6PC2*) is expressed in human α-cells in GWAS-allele specific manner

Bulk ATAC-seq analyses of sorted β-cells and single-nuclei chromatin accessibility profiling of sorted human islet cells (snATAC-seq) confirmed accessibility at the *G6PC2* promoter and nearby enhancers of β-cells (Fig. 1A), consistent with previous findings by O’Brien and colleagues illustrating *G6PC2* protein expression in β-cells(*12*) and numerous transcriptome analyses demonstrating *G6PC2* as one of the most highly expressed genes in β-cells (*24–27*). Importantly, our snATAC-seq and bulk ATAC-seq analyses revealed that the *G6PC2* promoter is also accessible in α-cells (Fig. 1A), which is further supported by single cell transcriptome data demonstrating expression of *G6PC2* in this cell type (*24–27*). Altogether, these findings suggest that *G6PC2* could play an important functional role in α-cell glucose-sensing, similar to its role in β-cells. To validate these findings, we performed immunofluorescent staining on non-diabetic human pancreatic sections for *G6PC2*, insulin, and glucagon, and found that the *G6PC2* protein is indeed expressed in human α-cells (Fig. 1B).

**Fig. 1:**
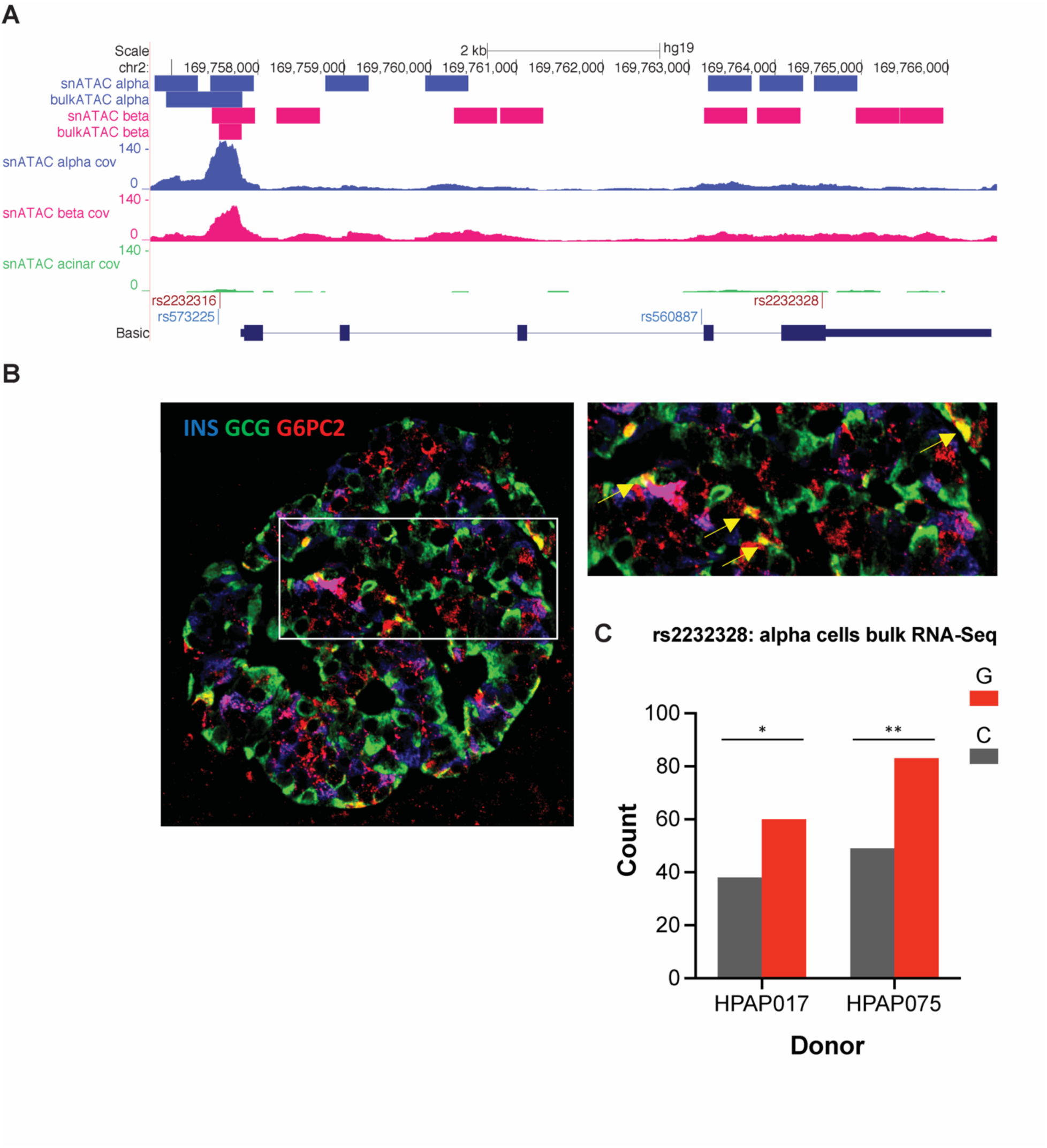
The islet-specific glucose-6-phosphatase catalytic subunit (G6PC2) is expressed in human α-cells. (**A**) Snapshot of the *G6PC2* region from the UCSC Genome Browser (build hg19) showing the following tracks: reliable α and β cell peaks from bulkATAC-seq; snATAC-seq signal for α, β, and acinar cells, and fasting plasma glucose (FPG) relevant SNPs. SNPs in high LD with each other (based on r^2^ in EUR) are shown with the same color. **(B)** Immunostaining of human non-diabetic pancreatic sections for G6PC2: G6PC2 is detected in a subset of α cells. **(C)** Allele-specific gene expression in α cells from two deceased European ancestry organ donors heterozygous for the reference (C) and alternate (G) alleles of rs2232328 from bulk RNA-seq data. P-values are from binomial tests: *p<0.05, **p<0.01.

The open chromatin region shared by α– and β-cells in the promoter of *G6PC2* spans two SNPs that are in linkage disequilibrium (LD) with SNPs associated with variation in fasted and fed blood glucose levels in people of European ancestry (EUR). Thus, rs573225 is found in LD with the lead SNP rs560887 for fasting glucose levels(*28*) (EUR; LD; r^2^>0.9 from HaploRegv4.1)(*29*) and rs2232316 is found in LD with rs2232328 (EUR; LD; r^2^>0.9 from HaploRegv4.1), an exonic missense variant (S342C) associated with variations in fasted and fed blood glucose(*30*). Both SNPs in the open chromatin region, rs573225 and rs2232316, were found to change FOXA binding motifs and are associated with altered *G6PC2* promoter activity in transfection assays(*15, 31*). To determine whether variants linked to fasting blood glucose are associated with *G6PC2* transcript levels in α-cells, we leveraged bulk RNAseq data from HPAP (https://hpap.pmacs.upenn.edu) to perform allele-specific expression analyses on the exonic rs2232328 SNP, for which the alternate allele is associated with an increase in *G6PC2* promoter activity *in* vitro(*15*). As mentioned above, increased expression of *G6PC2* in α-cells would be expected to result in increased glucagon secretion and thus increased fasting blood glucose levels. We had informative data for α-cells from two EUR donors and indeed identified statistically significant allelic imbalance, with increased expression of the alternate allele (G) – linked to higher fasting glucose – relative to the reference allele (C) in both donors (Fig. 1C). These data further support a functional role for *G6PC2* in controlling the glucose sensing set point in α-cells.

### Efficient ablation of *G6PC2* in α-cells of *G6pc2*^loxP/loxP^; Gcg-CreERT2 mice

To directly test the role of *G6pc2* in α-cell function, we derived a new mouse line in which the first three exons of *G6pc2* are flanked by *loxP* sites to enable cell type-specific gene ablation (Fig. 2A). These mice were bred to Gcg-CreERT2 mice(*32*) to allow for inducible, α-cell specific ablation of *G6pc2*. For our experimental model, we employed mice homozygous for the *G6pc2^loxP^* allele and hemizygous for the Cre allele (*G6pc2*^loxP/loxP^; Gcg-CreERT2; henceforth referred to as “α-G6PC2KO”). Our control group included mice homozygous for the *G6pc2*^loxP^ allele but lacking the Cre transgene, as well as mice carrying only the *Gcg-CreER^T2^* allele. Following tamoxifen administration, *G6pc2* expression was ablated in 88% of α-cells in αG6PC2KO mice compared to control mice, assessed by quantification of glucagon positive/ *G6pc2* negative cells in immune-stained pancreatic sections and consistent with the recombination efficiency reported previously for the Gcg-CreERT2 line(*32*) (Fig. 2B-C). To exclude the possibility that changes in islet cell number or morphology contribute to any potential alterations in α-cell function in α-G6PC2KO animals, we assessed the proportion of α– and β-cells out of all endocrine cells in the islet by co-staining for the pan-endocrine marker Chromogranin A (CgA) and quantified α– and β-cell proliferation using ethynyl deoxyuridine (EdU) labeling(*11*). We observed no changes in the proliferation rate of α– and β, nor a significant change in their relative proportion in islets of 12-week-old α-G6PC2KO versus age-matched control mice (Supplementary Fig. S1A-C). Additionally, the number and distribution of glucagon^+^CgA^+^ and insulin^+^CgA^+^ cells were similar between α-G6PC2KO and control islets, demonstrating that α-cell specific ablation of *G6pc2* did not impact islet composition (Supplementary Fig. S1D-F).

**Fig. 2:**
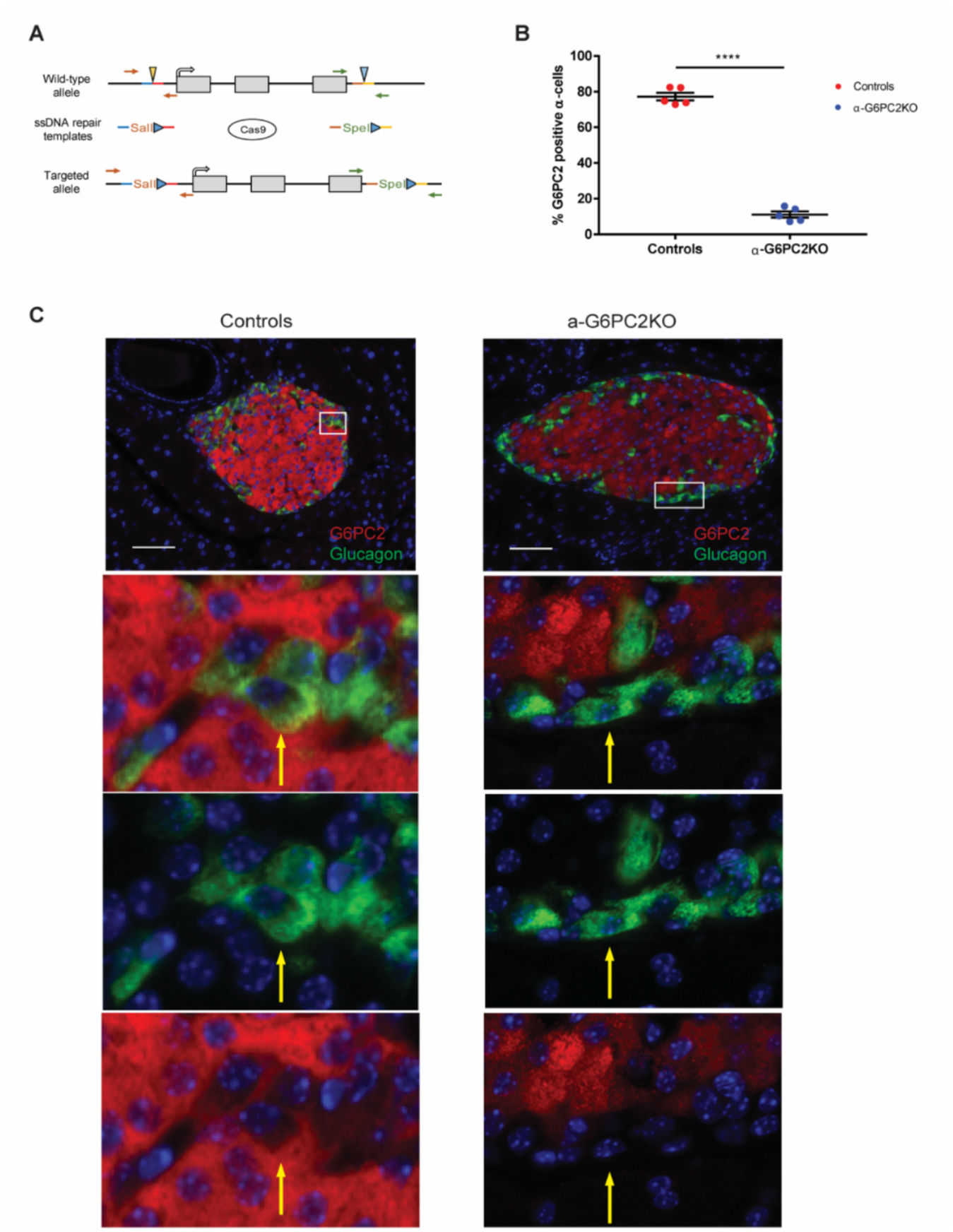
Efficient ablation of *G6pc2* in Gcg^+^ cells in G6pc2^loxP/loxP^; Gcg-CreERT2 mice. **(A)** Design of conditional null mutation in *G6pc2*: (top) *G6pc2* gene structure, with triangles indicating binding sites for gRNAs used for Cas9-assisted gene targeting, (middle) repair templates for introduction of the 5’ and 3’ loxP sites, where loxP sites (blue arrowhead) are marked by SalI or SpeI restriction endonuclease sites, (bottom) Targeted conditional null allele, with PCR primers used for genotyping (arrows). **(B)** Ablation efficiency of *G6pc2* from Gcg^+^ cells in the α-G6PC2KO model. **(C)** Representative immunofluorescent staining of G6pc2 and Glucagon in the pancreas from α-G6PC2KO and control mice. P-values from unpaired Student’s t-test: ****p<0.0001.

### Glucose cycling in islets from α-G6PC2KO mice

Previous studies using germline *G6pc2* null mice have implicated a role for G6PC2 in glucose cycling(*12, 14*). To assess the functional effect of our genetic manipulation, we performed *ex vivo* glucose uptake studies. Islets from α-G6PC2KO and control mice were incubated with 2– NBDG – a fluorescently labeled 2-deoxyglucose analog – and fluorescence measurements were collected post-washout with glucose-free media. 2-deoxyglucose can enter the cell as efficiently as glucose and is phosphorylated by GCK or other hexokinases; however, it cannot be further metabolized in the glycolytic pathway. When de-phosphorylated by G6PC2, 2-deoxyglucose can exit the cell again via the facilitative glucose transporters (GLUT2 and GLUT1) located in the plasma membrane. We observed that the 2-NBDG signal was similar between α-G6PC2KO and control islets immediately post-washout (T=0 min), suggesting no significant differences in basal glucose uptake (Fig. 3A). Remarkably, assessment of fluorescence following washout at 5 min and 15 min revealed a noticeably higher 2-NBDG signal in α-G6PC2KO (∼18% of baseline) compared to control islets (∼2% of baseline) (Fig. 3B-C). Given the findings from our islet composition studies, where we confirmed that α-cells comprise approximately 20% of murine islet cells (Supplementary Fig. S1E), in addition to the previously noted efficient recombination in the α-G6PC2KO model (Fig. 2B-C), we suggest that these glucose uptake results can be explained by the absence of futile cycling in G6PC2-deficient α-cells, which ‘traps’ 2-NBDG inside the mutant α-cells.

**Fig. 3:**
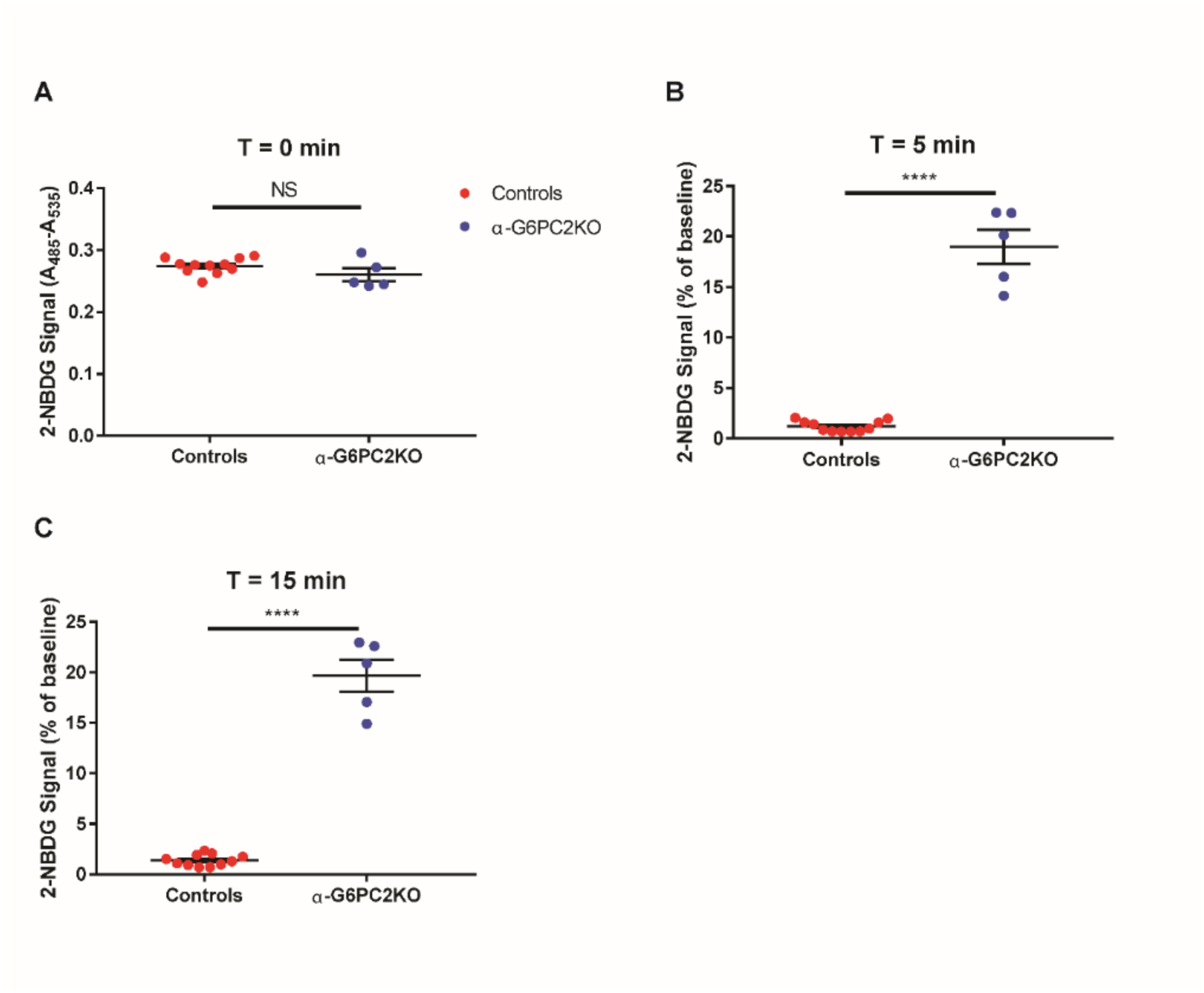
Higher levels of 2-NBDG fluorescence in pancreatic islets from G6pc2^loxP/loxP^; Gcg– CreER^T2^ mice. Isolated pancreatic islets were incubated with 2-NBDG for 30 minutes, and 2– NBDG fluorescence was measured at **(A)** 0 minutes, **(B)** 5 minutes, and **(C)** 15 minutes post–washout with glucose-free media. (n=5 mice for α-G6PC2KO, n=11 mice total for *G6pc2^loxP/loxP^* and *Gcg-CreER^T2^*). P-values from unpaired Student’s t-test: ****p<0.0001.

### α-cell *G6pc2* ablation alters glucose homeostasis

We employed cohorts of both male (Fig. 4) and female (Supplementary Fig. S2) mice for our phenotyping studies. Weekly measurements of body weight and *ad libitum* blood glucose starting at 4 weeks of age revealed no significant differences between α-G6PC2KO and control mice pre– and post-tamoxifen administration (Fig. 4A-B and Supplementary Fig. S2A-B). Intraperitoneal (i.p.) glucose tolerance tests (IPGTT) conducted 2-3 weeks after the final tamoxifen injection revealed no differences between α-G6PC2KO and control mice (Fig. 4C-D and Supplementary Fig. S2C-D). However, an assessment of glucagonemia before glucose administration and 5 minutes after, when glucose levels and thus suppression of glucagon secretion peak, revealed lower glucagon levels in both male and female α-G6PC2KO mice relative to controls (Fig. 4E and Supplementary Fig. S2E), as expected from increased α-cell glycolytic flux in the absence of futile glucose cycling. The fasting and glucose-suppressed glucagon levels were reduced in the absence of any significant difference in insulin levels between α-G6PC2KO and control mice (Fig. 4F and Supplementary Fig. S2F) excluding the possibility that a paracrine inhibitory effect of insulin is indirectly causing the reduction in glucagon secretion.

**Fig. 4:**
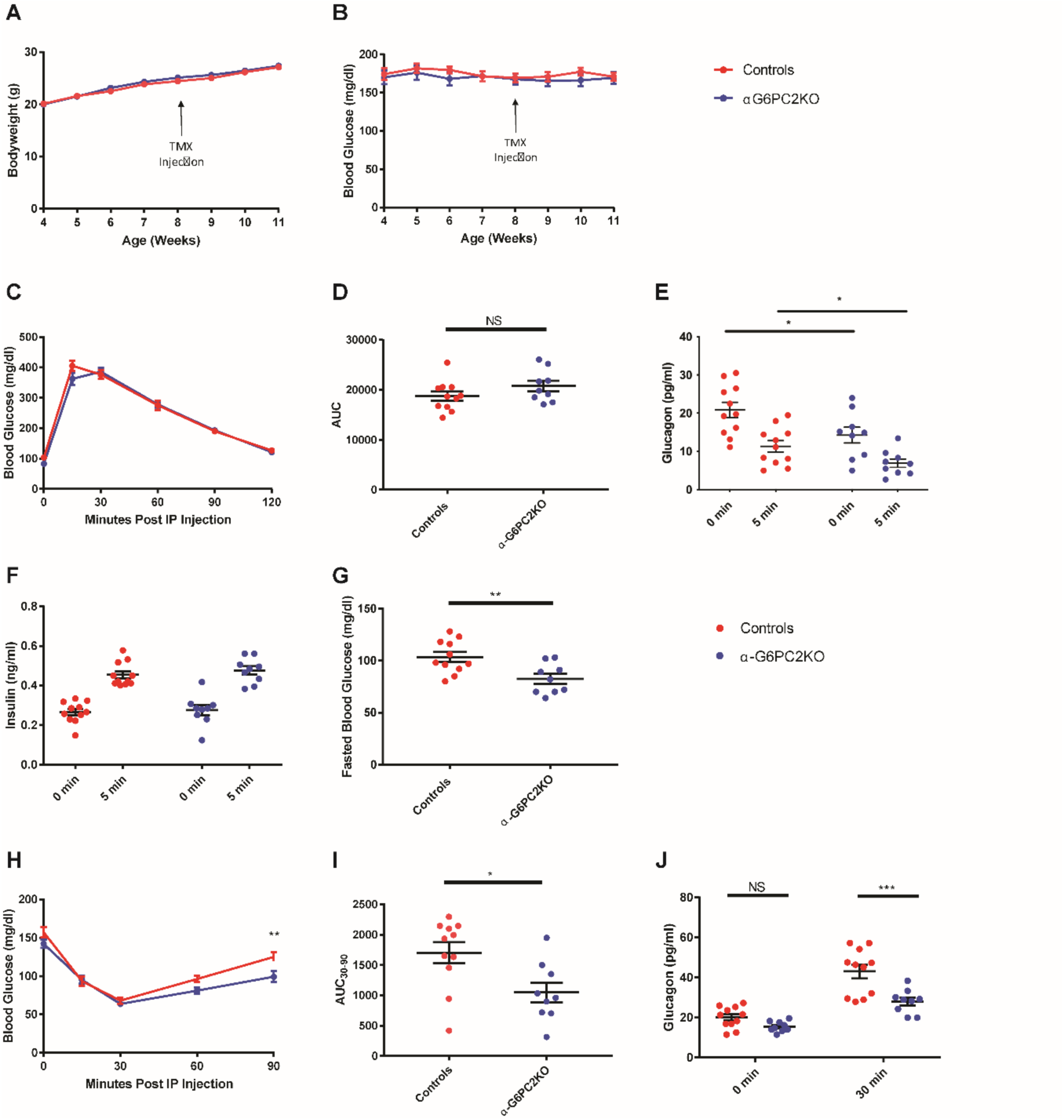
α-cell G6PC2 ablation alters glucose homeostasis in adult male mice. (**A**) Body weights of male α-G6PC2KO and control mice (n=9 for α-G6PC2KO, n=11 total for *G6pc2^loxP/loxP^*and *Gcg-CreER^T2^)*. **(B)** *Ad libitum* blood glucose for male α-G6PC2KO and control mice (n=9 for α-G6PC2KO, n=11 total for *G6pc2^loxP/loxP^* and *Gcg-CreER^T2^)*. **(C)** Intraperitoneal glucose tolerance test (1mg/g bodyweight) (n=9 for α-G6PC2KO, n=11 total for *G6pc2^loxP/loxP^* and *Gcg-CreER^T2^)*. **(D)** Area Under the Curve (AUC) from intraperitoneal glucose tolerance test in (C). **(E)** Plasma glucagon and **(F)** plasma insulin in mice fasted for 16 hours or 5 min after glucose injection in (**C**). **(G)** 16 hour fasted blood glucose levels (n=9 for α-G6PC2KO, n=11 total for *G6pc2^loxP/loxP^*and *Gcg-CreER^T2^*). **(H)** Intraperitoneal insulin tolerance test (0.75U/kg bodyweight) (n=9 for α– G6PC2KO, n=11 total for *G6pc2^loxP/loxP^* and *Gcg-CreER^T2^*). **(I)** Area Under the Curve between 30– and 90-min post-insulin injection (AUC_30-90_) from intraperitoneal insulin tolerance test in (H). **(J)** Plasma glucagon in mice fasted for 4 hours or 30 min after insulin injection in (H). (*p < 0.05, **p < 0.01, ***p < 0.001 vs. genetically unmodified control). P-values from unpaired Student’s t-test or two-way ANOVA with *post hoc* Bonferroni test: *p < 0.05, **p < 0.01, ***p < 0.001.

Overnight fasting of 16 hours prior to the IPGTT resulted in a significantly lower blood glucose levels in male α-G6PC2KO mice relative to control mice (Fig. 4G). Although we did not detect a statistically significant difference between female α-G6PC2KO and control mice in response to the 16-hour fast, we observed a trend of reduced blood glucose and glucagon levels in female α-G6PC2KO mice when fasting was prolonged to 24 and 36 hours while insulin levels remained similar to control mice (Supplementary Fig. S2G and Supplementary Fig. S3A-C).

Next, we performed an i.p. insulin tolerance test (ipITT) to further interrogate the effect of hypoglucagonemia in α-G6PC2KO mice on the response of the periphery to hypoglycemia, while not anticipating changes in liver insulin sensitivity. Notably, male and female α-G6PC2KO mice recovered more slowly from insulin-induced hypoglycemia relative to control mice, which was confirmed by AUC analysis between the 30– and 90-minute timepoints post-insulin administration (Fig. 4H-I and Supplementary Fig. S2H-I), concomitant with significant hypoglucagonemia at 30 minutes post-injection (Fig. 4J and Supplementary Fig. S2J). Again, these data are consistent with an altered setpoint for glucose suppression of glucagon secretion.

Given the proposed role of insulin as a paracrine inhibitor of glucagon release(*11*), and although insulin levels were similar between G6PC2KO and control mice, we sought to further rule out the possibility that the reduction in glucagon levels was a result of increased α-cell insulin sensitivity. Thus, we treated mice with either the SGLT1 inhibitor phloridzin, which lowers blood glucose by preventing renal glucose reabsorption, or vehicle control for 14 days to examine the effect of hypoglycemia on glucagon secretion in α-G6PC2KO mice independent of insulin(*33*). Both phloridzin-treated groups exhibited a decrease in blood glucose level as expected, with noticeably lower levels in the α-G6PC2KO mice after 14 days of treatment (Fig. 5A-B). Furthermore, α-G6PC2KO mice exhibited significantly lower glucagon levels at 7 and 14 days when compared to controls, while no change was detected in insulin levels, confirming that our observations from the ipITT were due to changes in α-cell intrinsic glucose sensing, rather than alterations in α-cell insulin sensitivity (Fig. 5C-D).

**Fig. 5:**
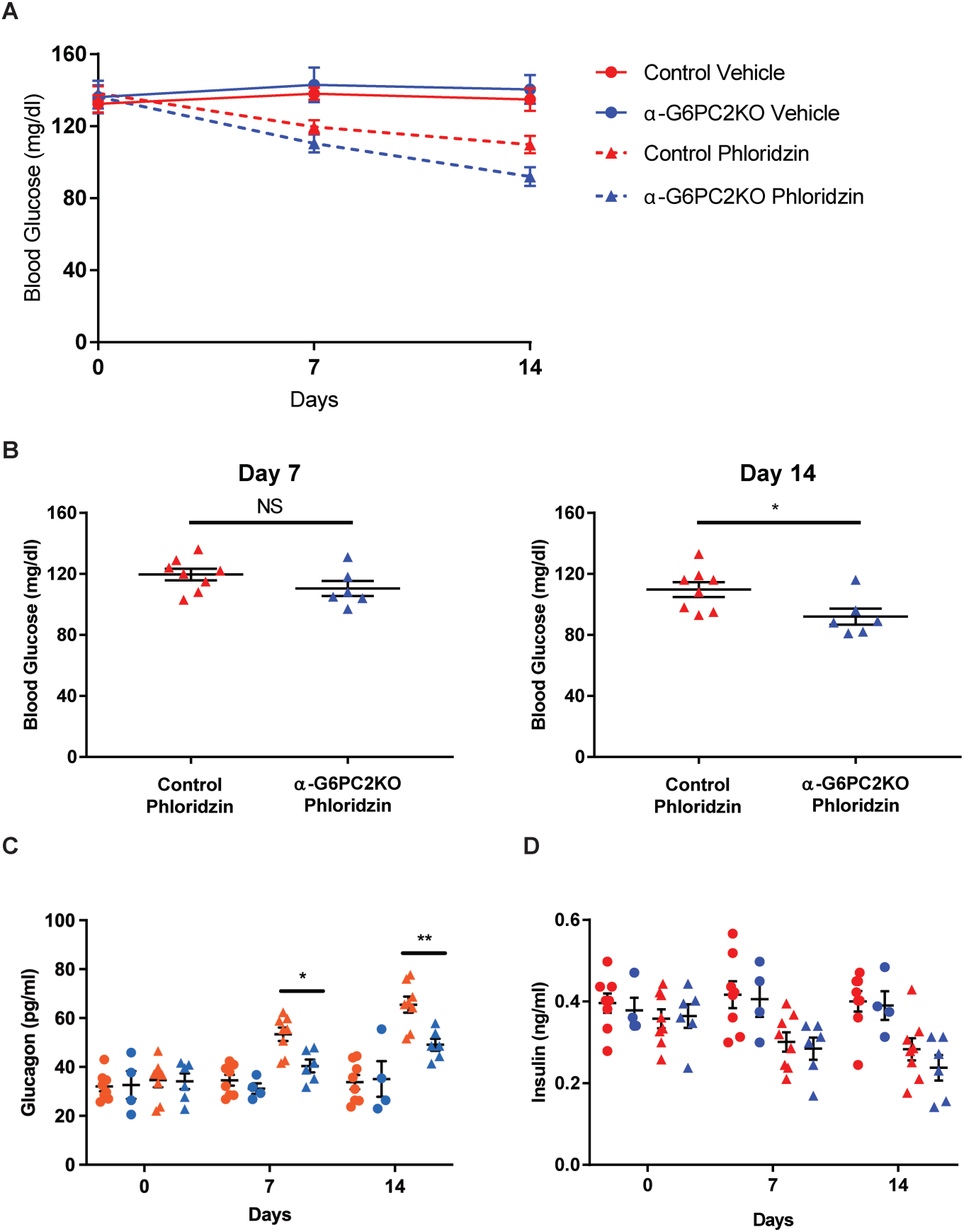
Ablation of α-cell G6PC2 impairs the counter-regulatory response to phloridzin-induced hypoglycemia. **(A)** *Ad libitum* blood glucose for female α-G6PC2KO and control mice (n=8 for Control Vehicle, n=4 for α-G6PC2KO Vehicle, n=8 for Control Phloridzin, n=6 for α– G6PC2KO Phloridzin) monitored on days 0, 7, and 14. **(B)** *Ad libitum* blood glucose for phloridzin-treated α-G6PC2KO and control mice on days 7 and 14 in (A) (n=5 for Control Phloridzin, n=4 for α-G6PC2KO Phloridzin). **(C)** Plasma glucagon and **(D)** insulin levels were assessed on days 0, 7, and 14 for the phloridzin-induced hypoglycemia experiment in (**A**). P-values from unpaired Student’s t-test or two-way ANOVA with *post hoc* Bonferroni test: *p<0.05.

### Ablation of *G6pc2* in α-cells enhances glucose-suppression of glucagon secretion (GSGS*) ex vivo*

The fine-tuned control of glucagon secretion is a complex process involving numerous paracrine, hormonal and neuronal signals acting under different physiological conditions in a stimulatory and inhibitory manner(*34*). Therefore, to isolate the intrinsic effects of *G6pc2* ablation on glucagon secretion from the possible impact of circulatory factors and innervation, we conducted *ex vivo* static batch incubation experiments on isolated islets from α-G6PC2KO and control mice (*11*). Islets were incubated in a medium containing a physiological 4mM amino acid mixture to stimulate glucagon release, with progressively increasing concentrations of glucose added to suppress glucagon secretion. Finally, islet cells were depolarized with 30mM KCl to assess the readily releasable pool of secretory granules, before being lysed for hormone content analysis. Incubation of islets in 4mM amino acids in the absence of glucose stimulated glucagon secretion similarly in α-G6PC2KO and control islets (Fig. 6A for islets isolated from male mice and Supplementary Fig. S4A for islets isolated from female mice). However, when incubated in the presence of 0.5mM, 1mM, 3mM, or 5mM glucose, α-G6PC2KO islets secreted significantly less glucagon relative to control islets, confirming a left shift in the GSGS curve (Fig 6A, Supplementary Fig. S4A). Next, we measured insulin secretion levels from the same islets to rule out the possibility that the observed inhibition in glucagon secretion was a secondary consequence of altered insulin levels within the islet. As shown in Fig. 6C-D and Supplementary Fig. S4C-D, the secretion of insulin in response to glucose, as well as insulin content, were similar in α-G6PC2KO and control islets.

**Fig. 6:**
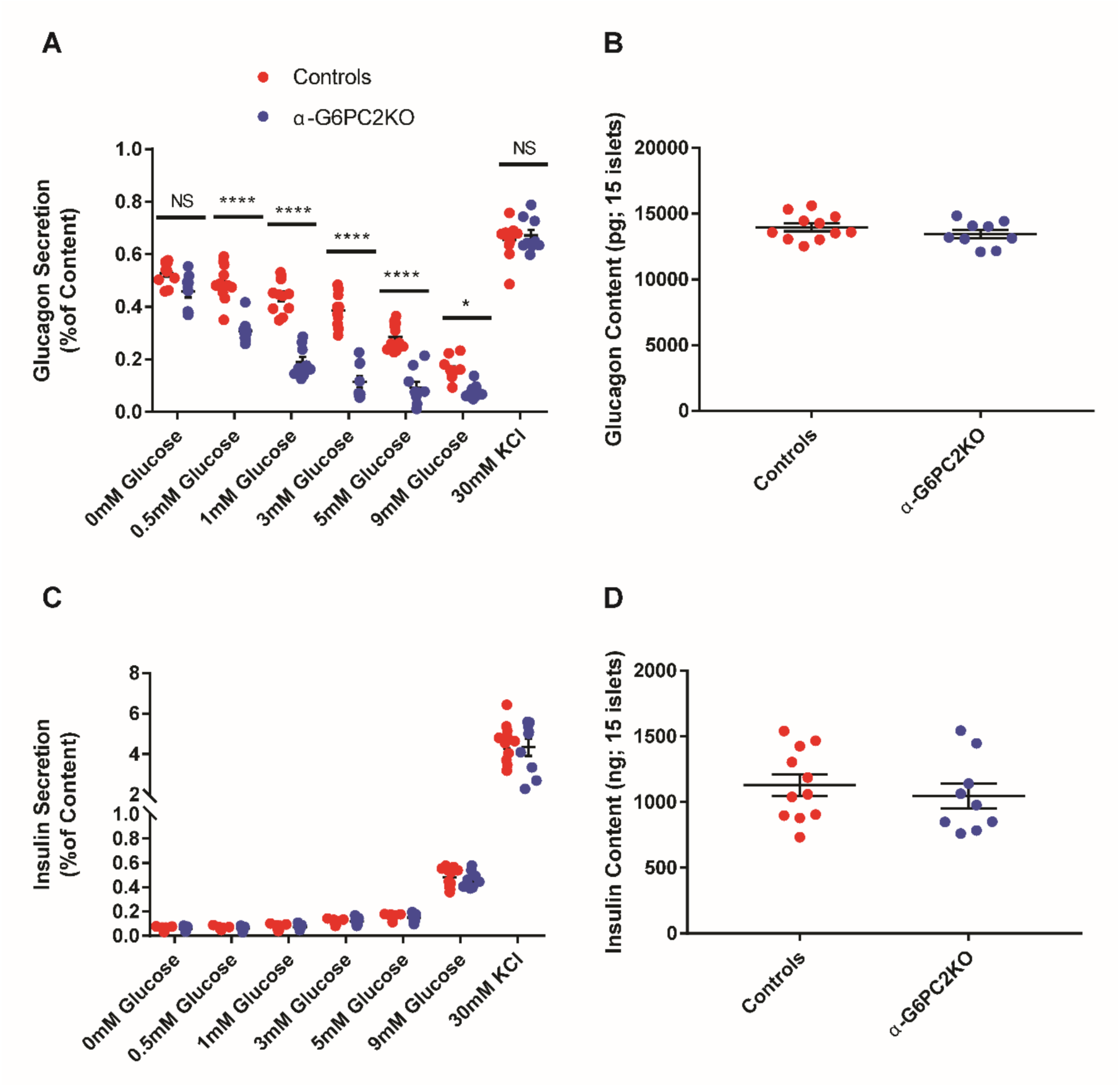
Glucose is more potent in suppressing glucagon secretion in isolated pancreatic islets from male α-G6PC2KO mice. **(A)** Glucagon secretion and **(B)** content from isolated pancreatic islets in the presence of the indicated glucose concentrations with 4mM amino acid mixture or 30mM KCl (n=9 for α-G6PC2KO, n=11 total for *G6pc2^loxP/loxP^*and *Gcg-CreER^T2^*). **(C)** Insulin secretion and **(D)** content from isolated pancreatic islets in the presence of the indicated glucose concentrations with 4mM amino acid mixture or 30mM KCl (n=9 for α-G6PC2KO, n=11 total for *G6pc2^loxP/loxP^* and *Gcg-CreER^T2^)*. P-values from two-way ANOVA with *post hoc* Bonferroni test:*p < 0.05, ****p < 0.0001.

### shRNA-mediated ablation of *G6PC2* limits glucagon release in α-cell-only human pseudo-**islets**

To confirm that the observations in mouse islets are also relevant to human α-cells, particularly considering species-specific differences in islet function such as the set-point for normoglycemia, we transduced sorted human α-cells obtained from non-diabetic deceased organ donors with lentiviral particles expressing shRNA against *G6PC2* (shG6PC2) or a non-silencing control (NS) and allowed them to aggregate for 4 days to form pseudo-islets while expressing the shRNA transgene (Fig. 7A). *G6PC2* mRNA levels were reduced by 80% as determined by qPCR (data not shown). To test the effect of downregulation of *G6PC2* in human α-cells on glucagon secretion, we measured GSGS using a static batch incubation assay in a similar setting to the *ex vivo* mouse experiment (see above and Methods). As shown in Fig. 7B, *G6PC2* ablation enhanced glucose inhibition of glucagon secretion, resulting in a left shift in the GSGS curve, confirming that *G6PC2* performs similar functions in human and mouse α-cells. Total glucagon content was not significantly different between the two experimental groups.

**Fig. 7:**
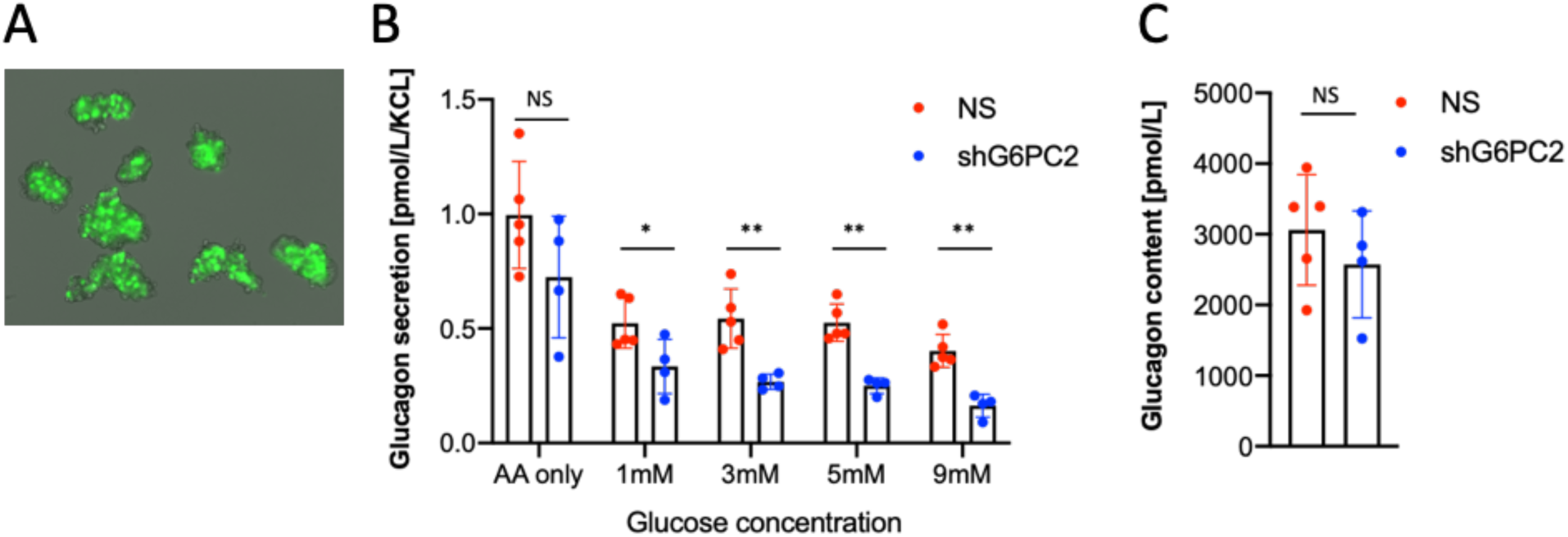
shRNA-mediated ablation of G6PC2 limits glucagon release in α-cell enriched human pseudo-islets. **(A)** Human α-cell pseudo-islets transduced with lentiviral particles expressing shRNA against *G6PC2* (shG6PC2) and GFP, 20X magnification. **(B)** Representative graph showing glucagon secretion assay on pseudo-islets transduced with lentiviral particles expressing shRNA against G6PC2 and GFP (G6PC2, blue) or a non-silencing control (NS, red), 12 pseudo-islets per replicate, 4-5 replicates per condition. **(C)** Glucagon content. P-values was determined by unpaired Student’s t-test. *, p< 0.05, **, p<0.01, ***, p<0.001, NS, not-significant.

## Discussion

Here we have shown that glucagon secretion by pancreatic α-cells is regulated by glucose not only via the forward glycolytic reaction catalyzed by glucokinase(*9–11*), but also by the reverse reaction dependent on *G6PC2*, which creates a futile cycle by converting glucose-6-phosphate to glucose and inorganic phosphate. Using inducible, cell type-specific gene ablation of *G6pc2* in mice we document a dramatic shift in the set-point for glucose suppression of glucagon secretion, resulting in more than half-maximal reduction in glucagon release from isolated islets at 1 mM glucose (Fig. 6). This effect results in lower fasting glucose levels (Fig. 4G) as the mutant α-cells cannot mount a strong glucagon response. Thus, *G6PC2* is a critical regulator of the setpoint for glucose suppression of glucagon secretion in murine pancreatic α-cells.

Extensive evidence confirms the relevance of these findings in humans. Polymorphisms in the *G6PC2* gene are significantly associated with alterations in fasting blood glucose and HbA1c levels(*15–18*) and were previously studied for their potential impact on *G6PC2* transcription and splicing in β-cell lines(*19*). Here we showed that relevant polymorphisms effect α-cell *G6PC2* chromatin accessibility (ATACseq data, Fig. 1A), and gene expression (Fig. 1C), and, finally, that decreased *G6CP2* expression in human α-cells results in a leftward shift in the glucose inhibition of glucagon secretion (Fig. 7), similar to that observed in α-cell specific ablation of *G6PC2* expression in intact mouse islets.

As established decades ago, glucokinase activity, catalyzing the phosphorylation of glucose to glucose-6-phosphate (G6P), is the key determinant for regulating hormone secretion in glucose sensing cells, coupling variations in glucose levels to hormone output(*35, 36*). When compared to other hexokinases, glucokinase possesses unique kinetic properties, including its sigmoidal substrate dependency curve – with an inflection point at a physiologically relevant glucose concentration – and positive cooperativity with glucose binding (Hill coefficient of 1.7) (Fig. 8A). In contrast, cells that do not serve as glucose sensors express hexokinase I – III, which have a high affinity for glucose, rendering it insensitive to changes in glucose concentrations within the physiological range (Fig. 8B). As first suggested by O’Brien and colleagues for insulin secretion in β-cells(*12, 13*), our model, based on the findings reported above, proposes that *G6PC2* fine-tunes glucose sensing in α-cells; at low G6P concentrations, the presence of *G6PC2* further decreases glycolytic flux even below that determined by the sigmoidal glucose response curve of GCK alone, thus enabling tighter control over glucagon secretion at low blood glucose levels (Fig. 8C). However, once G6P concentrations rise, the limited catalytic capacity of *G6PC2* is overwhelmed, and glycolytic flux is largely determined by GCK.

**Fig. 8:**
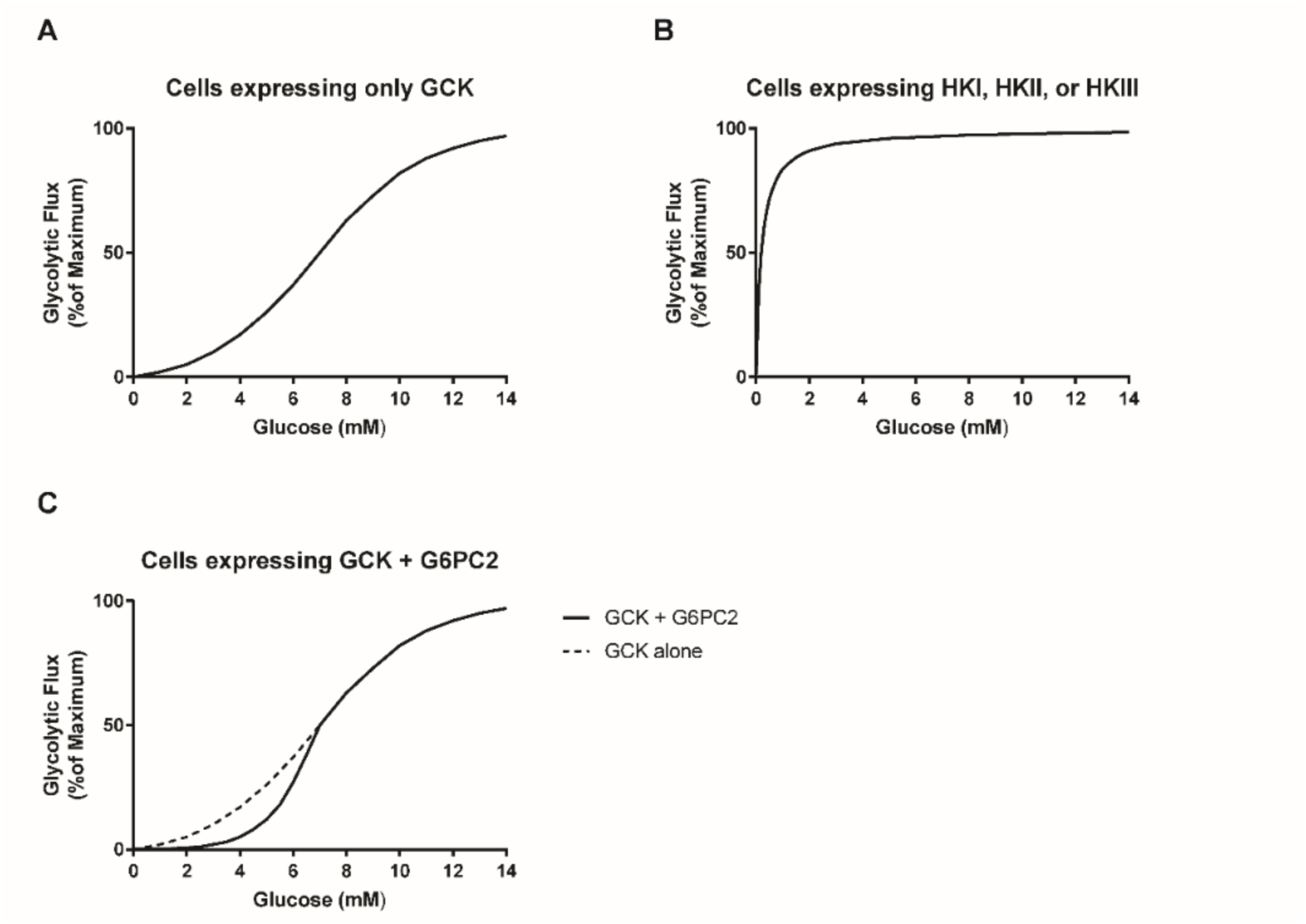
The glucokinase/G6PC2 futile substrate cycle regulates glycolytic flux. **(A)** Schematic illustrating the effect of glucokinase on glycolytic flux in the absence of G6PC2 activity. Glucokinase possesses unique kinetic properties, such as the depicted sigmoidal substrate dependency curve, that allows it to couple variations in glucose levels to changes in glycolytic flux and hormone output. **(B)** Schematic illustrating the effect of hexokinase (HK) I, II, and III – which are expressed in cells that do not serve as glucose sensors – on glycolytix flux. HK I, II, and III possess a high affinity for glucose, rendering them insensitive to changes in glucose concentration within the physiological range. **(C)** Schematic illustrating the effect of the glucokinase/G6PC2 futile substrate cycle on glycolytic flux. The model proposes that at low glucose-6-phosphate concentrations, the presence of *G6PC2* ensures very low glycolytic flux, even lower than enabled by the sigmoidal glucose response curve of GCK alone. However, once glucose-6-phosphate concentrations rise, the catalytic capacity of G6PC2 is overwhelmed, and glycolytic flux is largely determined by GCK.

Discovering safe and effective pharmacological treatments for type 2 diabetes patients remains a significant challenge(*37*). Due to the critical role of glucokinase in maintaining blood glucose homeostasis, several allosteric glucokinase activators (GKAs) have been evaluated for the treatment of Type 2 diabetes patients(*38–41*), but most were not developed further due to side effects such as hyperlipidemia and hypertension caused by elevated GCK activity in hepatocytes(*42–44*). In principle, inhibitors specific to G6PC2, but not targeting the liver isoform G6PC1, will increase glycolytic flux in both β– and α-cells and thus affect islet hormones in a bidirectional fashion, increasing insulin and lowering glucagon output. Because the *G6PC2* isoform is exclusively expressed in islet cells, side effects in hepatocytes are not expected, making it an attractive drug target. Since *G6PC2* has evolutionarily diverged substantially from the liver enzyme encoded by *G6PC1*(*45, 46*), the distinct properties and structural divergence of the two proteins may facilitate the identification of small molecule inhibitors specific for G6PC2. In evidence of this, Petrolonis and colleagues previously conducted high-throughput screening of a small molecule library and identified several compounds that inhibited G6PC2 enzymatic activity without affecting G6PC1 *in vitro*(*47*). Thus, G6PC2 inhibitors that can act through a novel, bi– hormonal mechanism should be developed and tested further as potential treatment for patients with Type 2 diabetes.

## Methods

### Mice

For the derivation of the *G6PC2*^LoxP/LoxP^ allele, we employed Crispr/Cas9 assisted homologous recombination in zygotes from C57BL/6 mice. Two guide RNAs, targeting sequences upstream of exon 1 and downstream of exon 3 were employed, together with two repair templates that introduced loxP sites together with a SalI restriction site at the 5’ location, and a SpeI restriction site at the 3’ location. Microinjection was performed by Penn’s Transgenic and Chimeric Mouse Facility. Offspring were screened by PCR followed by restriction endonuclease digest. The derivation of the *Gcg-CreER^T2^*allele has been reported previously(*32*). α-G6PC2KO mice were homozygous for the *G6PC2*^LoxP/LoxP^ allele and heterozygous for the *Gcg-CreER^T2^* allele. All mice were maintained on a C57BL/6 background and housed on a standard 12-h light/12-h dark cycle with *ad libitum* access to food and water. Bodyweight and random blood glucose levels were measured weekly beginning at 4 weeks of age. To induce Cre activity, tamoxifen (Sigma–Aldrich #T5648) was administered to 8-week-old mice at 50 μg/g bodyweight via 3 intraperitoneal (i.p.) injections at 24-h intervals. *In vivo* metabolic studies were conducted two weeks following the final tamoxifen injection to ensure sufficient α-cell-specific genetic modification(*11, 32*). All animal procedures were approved by the University of Pennsylvania Institutional Animal Care and Use Committee.

### Glucose tolerance and insulin tolerance tests

Glucose tolerance tests and insulin tolerance tests were performed according to previously published protocols(*11*). For glucose tolerance tests, mice underwent a 16-h overnight fast and then received an i.p. injection of 1 mg/g bodyweight of D-glucose in sterile phosphate buffered saline (PBS). Measurements of blood glucose were taken at 0-, 15-, 30-, 60-, 90-, and 120-min post-injection using an automatic glucometer (Zoetis). The area under the curve was calculated using GraphPad PRISM software (v.7.0), using the time zero glucose measurement as the baseline. For assessment of glucose-stimulated insulin secretion (GSIS) and glucose suppression of glucagon secretion (GSGS), blood was collected from the tail vein at 0– and 5-min post-injection and placed on dry ice with aprotinin (Sigma Aldrich #A6279; final concentration: 0.167 mg/ml) to prevent glucagon degradation. Plasma insulin and glucagon concentrations were quantified using enzyme-linked immunosorbent assay (ELISA, Crystal Chem #90080; Crystal Chem #81518).

For insulin tolerance tests, mice underwent a 4-h fast and then received an i.p. injection of 0.75 U/kg bodyweight of insulin. Measurements of blood glucose were taken at 0-, 15-, 30-, 60, and 90-min post-injection using an automatic glucometer (Zoetis). The area under the curve was calculated between 30 and 90 min using GraphPad PRISM software (v.7.0) to assess the counterregulatory response. To assess glucagonemia under conditions of insulin-induced hypoglycemia, blood was collected from the tail vein at 0– and 30-min post-injection, and plasma glucagon concentrations were quantified as described above.

### Fasting Experiments

Individually housed female mice underwent a 36-h fast, and blood glucose was measured every 12 h using an automatic glucometer (Zoetis). Blood was collected from the tail vein at each 12 h interval, and plasma insulin and glucagon concentrations were quantified using the methods described above.

### Phloridzin Treatment Experiments

Mice were treated with 0.4 mg/g bodyweight of phloridzin or vehicle twice a day for 2 weeks(*33*). Blood glucose was measured on days 0, 7, and 14 using an automatic glucometer (Zoetis). Additionally, blood was collected from the tail vein on Days 0, 7, and 14, and plasma insulin and glucagon concentrations were quantified using the methods described above.

### Islet isolation and culture

Mouse islets were isolated using published protocols(*11*). Islet isolation required inflation of the pancreas followed by digestion with collagenase (Roche #11213873001) at 37 °C. Islets were then enriched via density gradient centrifugation with Ficoll–Paque (GE 45-001-751) and underwent 3 rounds of hand-picking to separate islets from exocrine tissue. Islets were cultured for three days prior to static batch incubations. The culture media consisted of RPMI 1640 (11 mM glucose; Thermo #21870076) supplemented with 10% fetal bovine serum (FBS), 2 mM glutamine, 1 mM sodium pyruvate (Thermo #11360070), 10 mM HEPES, and 1% antibiotic antimycotic reagent (ThermoFisher Scientific #15240096). The pH of the culture media was adjusted to 7.3–7.4.

### Static batch incubation in mouse islets

Static batch incubation experiments of mouse islets were performed and analyzed according to previously published protocols(*11*). Briefly, batches of 15 size-matched islets were preincubated in Krebs buffer containing 9 mM glucose for 1 h, washed and re-incubated with 9 mM glucose, followed by incubations with varying concentrations of glucose for a period of 30 min each, in the presence of a physiological 4 mM amino acid mixture. At the end of the experiment, islets were incubated with 30 mM KCl to confirm islet viability. After each incubation, islets were gently centrifuged at 230 rcf for 5 min, and supernatants removed and stored at −80 °C with aprotinin (Sigma Aldrich #A6279; final concentration: 0.025 mg/ml) to prevent glucagon degradation. Finally, islets were lysed using NP-40 lysis buffer (0.005% NP-40; pH 8.8) for total hormone content analysis. Static batch incubation studies were performed in triplicate. Insulin and glucagon concentrations were quantified using ELISA (Crystal Chem #90080; Crystal Chem #81518).

## 2-NBDG Glucose Uptake Assay

To measure glucose uptake, batches of 15 size-matched islets were incubated with 100 μg/mL 2-NBDG (ab235976), a fluorescently labeled 2-deoxyglucose analog, in glucose-free RPMI (Roswell Park Memorial Institute) medium for 30 min. After the incubation with 2-NBDG, islets were centrifuged at 400xg for 5 min, and the supernatant was replaced with glucose-free media. Fluorescence was measured post-washout at 0-, 5-, and 15-min with glucose-free media by microplate reader at excitation and emission wavelengths of 485 nm and 535 nm, respectively. 2-NBDG glucose uptake studies were performed in triplicate.

### Human α-cell sorting, transduction and aggregation into pseudo-islets

Human islets from non-diabetic (ND) donors (Table S1) were obtained from PRODO laboratories and cultured overnight upon arrival at 37⁰C in Prodo Islet Media – Standard PIM(S) medium supplemented with 5% Human AB blood type serum PIM(ABS), Glutathione mixture PIM(G) and antibiotics mix PIM(3X). Islets were dissociated with accutase (C-41310, Sigma-Aldrich, 500μl for 500 islets) for 10 minutes at 37⁰C followed by quenching in 20% fetal bovine serum in PBS (Biological industries, Israel). Cells were washed with PBS and stained for fluorescence-activated cell sorting (FACS) with Anti-HPa3, clone HIC3-2D12, (MABS1998, Sigma-Aldrich) to enrich for α-cells (*48*). After sorting, the α-cells fraction was transduced with either G6PC2-shRNA or Scramble-shRNA control (NS) lentiviruses (Vector Builder, VB221003-1034hsq and VB010000-0009mxc respectively) at a ratio of about 1:5 MOI in PIM(S) overnight. Transduced cells were washed and seeded in an agarose mold prepared in a MicroTissues® 3D Petri Dish® (Z764043, Sigma-Aldrich) for 3-4 days to form pseudo-islets and express the transgene.

### Static GSGS assay of human pseudo-islets

Glucagon secretion assay was performed in Krebs-Ringer buffer (KRBB) containing 114.4mM NaCl, 5mM KCl, 24mM NaHCO_3_, 1mM MgCl_2_, 2.2mM CaCl_2_, 10mM HEPES and 0.5% BSA, adjusted to pH 7.35. Pseudo-islets (12 per replicate, 4-5 replicates per condition) were pre–incubated in KRBB without glucose or amino acids for 1 hour, then this buffer was replaced with KRBB containing 4mM amino acid mix (glucagon induction) which was collected after 1 hour followed by 40 minutes incubations in 1mM, 3mM, 5mM, 9mM glucose and finally 30mM KCL, all in the presence of 4mM amino acid mix. At the end of each incubation period, the supernatant was collected, aprotinin (A6279, Sigma) was added to 1:100 and the supernatant was stored at – 80°C to avoid degradation of glucagon. Pseudo-islets were lysed in NP-40 lysis buffer (Thermo Fisher Scientific #FNN0021) for the determination of total glucagon content. Glucagon levels were measured using an ELISA kit (Mercodia 10-1271-01) and normalized to 30mM KCl representing readily secretable hormone granules. Statistical analyses were performed using a one-tailed Student’s t test. Data are presented as mean ± SE.

### Immunofluorescent analysis

Formalin-fixed paraffin-embedded (FFPE) pancreas sections from mice were stained to assess cell percentage and proliferation(*11*). Pancreata were dissected, fixed in 4% paraformaldehyde (PFA) overnight at 4°C, embedded in paraffin, and cut into 6-μm sections. To obtain the maximal footprint from each mouse pancreas, three consecutive sections spaced 250 μm apart from each other were stained. Slides were deparaffinized in xylene and rehydrated through a series of ethanol washes. Slides were subjected to antigen retrieval for 2 h in 10 mM citric acid buffer (pH 6.0). Ethynyl deoxyuridine (EdU) labeling was employed to detect cells that had entered or completed S-phase. EdU (Thermo # E10187) was supplied continuously for 7 days in the drinking water at a concentration of 1 mg/ml. EdU^+^ cells were labeled using the ClickiT EdU Alexa Fluor 647 imaging kit (Invitrogen #C10640) according to the manufacturer’s instructions. Sections were blocked using CAS-Block (Invitrogen #008120) and incubated overnight at 4°C with the following primary antibodies as appropriate: rabbit anti-insulin (1:500; proteintech), rabbit anti-glucagon (1:500; proteintech) or mouse anti-glucagon (1:10,000; Millipore Sigma), or mouse anti-chromogranin A (1:300; proteintech). Slides were washed in PBS and incubated with secondary antibodies for 2 hours at room temperature (mouse Cy2 1:200 and rabbit Cy3 1:200, Jackson ImmunoResearch). Hoechst 33342 (Thermo #H3570) was used to counterstain nuclei. α-cell proliferation was calculated as the number of EdU^+^, glucagon^+^ cells normalized to the total number of glucagon^+^ cells. α-cell percentage was calculated as the number of CgA^+^, glucagon^+^ cells divided by the number of CgA^+^ cells. These sections were also co-stained for insulin to assess β-cell proliferation and β-cell percentage using analogous methods. Staining was visualized using a Keyence BZ-X800 fluorescence microscope and images were analyzed using FIJI software.

Pancreatic cryo-sections were stained to validate G6PC2 ablation in mouse α-cells(*12*). Pancreata were dissected, fixed in 4% paraformaldehyde (PFA) for 1 hour at 4°C, washed three times with PBS, and placed in 30% sucrose in PBS overnight at 4°C. Pancreata were then immersed in optimal cutting temperature (OCT) reagent for 2 hours before being frozen on a dry ice block. Pancreatic cryo-sections were rehydrated in PBS for 10 min, permeabilized with 0.3% Triton X-100 for 30 min and washed three times with PBST. Sections were blocked using CASBlock (Invitrogen #008120) and incubated overnight at 4°C with the following primary antibodies: rabbit anti-G6PC2 (1:50; a kind and generous gift from Dr. Howard Davidson) or mouse anti-glucagon (1:10,000; Millipore Sigma). Slides were washed in PBS and incubated for 2-h at RT with secondary antibodies for 2 hours at room temperature (mouse Cy2 1:200 and rabbit Cy3 1:200, Jackson ImmunoResearch). Hoechst 33342 (Thermo #H3570) was used to counterstain nuclei. Recombination efficiency was calculated as the number of G6PC2^+^, glucagon^+^ cells normalized to the total number of glucagon^+^ cells. Staining was visualized and analyzed as described above.

Paraffin sections obtained from the JDRF Network for Pancreatic Organ Donors with diabetes (nPOD) were stained to quantify the prevalence of G6PC2 expressing α-cells in human islets from non-diabetic (ND) donors. Slides were deparaffinized in xylene, rehydrated in serial concentrations of alcohol (100, 90, and 80%) and double distilled water. Antigen retrieval was performed in 10mM citrate buffer (pH=6) using a pressure cooker (Biocare Medical). Sections were blocked with CAS-Block (Invitrogen) and incubated overnight with the following primary antibodies: guinea pig anti-insulin (1:5; DAKO), mouse anti-glucagon (1:200; Abcam) or rabbit anti-glucagon (1:200; Cell signaling technology) and rabbit anti-G6PC2 (1:100; ENCO). We employed DAPI (1:100; Sigma-Aldrich) to counterstain DNA. After an overnight incubation with primary antibodies, the slides were washed three times and incubated with secondary antibodies for 2 hours at room temperature (guinea pig Cy2 1:200, mouse Cy3 1:400 and rabbit Cy5 1:400, Jackson ImmunoResearch). Following staining, aqueous mounting medium was applied and covered with a coverslip. Images were taken on an Olympus confocal microscope at 40X magnification.

### Mouse data analyses

Data are presented as mean ± SEM and were analyzed using GraphPad Prism software (v.7.0). Statistical analysis was performed using an unpaired two-tailed Student’s t-test when two groups were compared, one-way analysis of variance (ANOVA) with *post hoc* Bonferroni test when more than two groups were compared, and two-way ANOVA with *post hoc* Bonferroni test when two conditions were involved. Values are considered statistically significant when *p* < 0.05.

### Single nucleus ATAC-seq

Single-nucleus ATAC-seq (snATAC-seq) results shown in this study were obtained from analyzing 14 adult non-diabetic deceased organ donors analyzed by the Human Pancreas Analysis program (HPAP; https://hpap.pmacs.upenn.edu)(*49, 50*). Donor islets were washed and approximately 200 islets handpicked and trypsinized to create single cell suspensions. Nuclei were then isolated according to the Nuclei Isolation for Single Cell ATAC Sequencing protocol (CG000169) from 10X Genomics. snATAC-seq libraries were then prepared with Chromium Next GEM Single Cell ATAC Reagent Kits v1.1 (10X, PN-1000175, PN-1000161), according to the User Guide (RevD). Briefly, single nucleus suspensions were transformed into snATAC-seq libraries through transposition, gel bead-in-emulsion (GEMs) generation using a microfluidic chip, pre-amplification, emulsion breaking, DNA amplification, and purification. 2,717+/-1,496 (min=358) single nuclei were captured per donor using 10X single-nuclei ATAC kit and converted to sequencing libraries. Samples were indexed using the Single Index Plate N Set A (10X, PN1000212). Library quality and concentration were determined using the Bioanalyzer High Sensitivity DNA kit (Agilent, PN-5067-4626). Samples were sequenced on Illumina sequencers using paired-end (50×49bp) reads to a mean depth of 71,000 +/– 56,000 reads per cell (min=23,000). Read pairs were aligned to the genome using cellranger-atac-2.0.0. Analysis of snATAC-seq was performed using ArchR v.1.0.1 according to the tutorial(*51*). Firstly, fragment files (output from 10x cellranger-atac counts) for each donor were loaded into ArchR using ‘createArrowFiles’. At the same time, a tiled matrix was calculated, which breaks down the genome into 500bp non-overlapping bins and the insertion counts for each bin were tabulated to represent each single nucleus. Then, nuclei with < 1,000 number of fragments or < 4 enrichment for transcriptional start sites were filtered out. Doublets were filtered out after calling ‘addDoubletScores’ and ‘filterDoublets’ with default parameters. Dimensional reduction was performed using ‘addIterativeLSI’ with useMatrix=“TileMatrix”, iterations=3, and resolution=0.2. Harmony was applied to correct for batch effects using the dimensional reduced representation from ‘addIterativeLSI’ as input(*52*). Then, ‘addClusters’ and ‘addUMAP’ were used to identify clusters and project the cells to two-dimensional UMAP space. Clusters were annotated based on accessibility at cell type marker genes. Open chromatin peaks were called with the default ArchR’s method.

Using the cell type annotation from the previous step, fragments associated with cells from each non-diabetic donor for whom we also had bulk ATAC-seq (HPAP035 and HPAP036) and each cluster (alpha, beta, delta, and acinar) were merged to generate pseudo-bulk tracks. The ethnicity of these two donors was European American (EUR). A bed file including the start, end, and chromosome information of each read was created for each donor and each cell type. Bam files were constructed using bedToBam v2.30.0 with genome size file provided as part of the GRCh38 reference – 2020-A-2.0.0 available on the 10x Genomics website(*53*). Bam files were indexed using samtools v1.11 index(*54*). BAMscale v0.0.5 was used to generate scaled coverage track, with parameter –t 4(*55*). Lastly, tracks of the same cell type were merged across donors using bigWigMerge v2 and the output was sorted and converted to a bw file using bedGraphToBigWig v2.8(*56*). These data were then lifted over to hg19 using CrossMap v.0.6.1(*57*).

### Bulk ATAC-seq

Bulk ATAC-seq was performed using previously published protocols on FACS-sorted human α-cell and β-cell samples(*58, 59*). Briefly, cells were resuspended in ATAC-seq resuspension buffer (RSB; 10M Tris-HCl, pH 7.5, 10mM NaCl, and 3mM MgCl_2_ in nuclease-free H_2_O and centrifuged at 8,000xg for 4 minutes at 4°C. The supernatant was aspirated, and cell pellets resuspended and incubated with 50μl lysis buffer (0.1% v/v NP-40, 0.1% v/v Tween-20, 0.1% v/v Digitonin in RSB) on ice for 3 minutes. Following lysis, 1ml wash buffer (0.1% v/v Tween-20 in RSB) was added and nuclei were centrifuged at 500xg for 10 minutes at 4°C. The supernatant was aspirated, and nuclei were resuspended and incubated with 50 μl transposition buffer (25 μl 2× TD buffer, 16.5 μl PBS, 0.1% v/v Tween-20, 0.1% v/v Digitonin, 2.5 μl Tn5 transposase [Tagment DNA Enzyme 1], and 5 μl water) at 37 °C for 25 minutes in a thermomixer at 1,000 r.p.m. Following the transposition, samples were purified using the Qiagen MinElute Reaction Cleanup Kit. Transposed DNA fragments were amplified using Ad1_noMX and Ad2.* barcoded primers for 5-10 cycles and purified using Agencourt AMPure XP beads (Beckman Coulter #A63881) to remove contaminating primer dimers. Library quality was assessed using the Agilent High Sensitivity DNA Kit (Agilent Technologies #5067-4626), and DNA concentration was assessed via the QuBit dsDNA high-sensitivity assay kit (Thermo #Q32854). Samples were sequenced on Illumina sequencers using paired-end reads >49 bases to a depth of at least 25 million reads, with a median depth of 40 million reads.

We utilized bulk ATAC-seq data from α and β cells of EUR non-diabetic donors from HPAP. For each cell type, we only used those donors passing the purity checks described in the next section; 8 donors for alpha cells (HPAP019, HPAP035, HPAP038, HPAP040, HPAP045, HPAP054, HPAP069, HPAP077) and 5 donors for beta cells (HPAP019, HPAP035, HPAP036, HPAP040, HPAP045). For each cell type and each individual, a set of ‘reliable’ peaks was generated as follows. Adapters in paired reads in the fastq files were trimmed with cutadapt v3.5(*60*). Trimmed reads were aligned to hg19 with bowtie2 v.2.3.4.1(*61*). The following were filtered out from the resulting bam files: unmapped reads, reads mapping to chrM, reads overlapping ENCODE blacklisted regions, secondary alignments, reads failing platform/vendor quality checks, reads not properly paired, and alignments with quality <10. Finally, duplicates were removed and read mates fixed. For these manipulations we leveraged samtools v1.11, bedtools 2.15.0, and picard v1.141(*53, 54*). Narrow peaks were called using macs2 v2.2.7.1. with a q-value threshold of 0.05(*62*). ‘Reliable’ peaks were defined as those overlapping by at least 20% with a peak from single cell ATAC-seq in cells from the same type (α or β). For each cell type, we finally took all pairwise overlaps of reliable peaks between the donors and merged the results to generate the consensus peaks plotted in Figure 1A.

### Bulk RNA-seq and allele-specific expression analyses

RNA was isolated from FACS-sorted human α-cell and β-cell samples using the Qiagen DNA/RNA AllPrep Micro kit (<250,000 cells; Qiagen #80284), Qiagen DNA/RNA AllPrep Mini kit (250,000-500,000 cells; Qiagen # 80204), or Qiagen DNA/RNA Universal AllPrep kit (>500,000 cells; #80224). Briefly, cells were pelleted, lysed using a buffer containing β-mercaptoethanol, and homogenized using a QIAshredder spin column. The homogenized lysate was transferred to an AllPrep DNA Mini spin column and centrifuged, and the flow-through containing RNA was incubated with proteinase K. The total RNA was then transferred to a RNeasy Mini spin column, incubated with DNase I, and then washed multiple times and eluted. Bulk RNA-seq libraries were then prepared using the Illumina TruSeq Stranded Total RNA Library Prep Gold kit (Illumina #20020599).

Bulk RNA-seq data from α and β cells of HPAP donors were first analyzed to assess purity. To this end, we employed MuSiC for cell type deconvolution(*63*), leveraging scRNA-seq data from 17 HPAP non-diabetic donors, with cells annotated as acinar, alpha, beta, delta, ductal, endothelial, mesenchymal, or PP. To perform deconvolution, we used a list of 244 pancreatic marker genes compiled from previous pancreatic scRNA-seq studies. Prior to deconvolution, samples with a total number of counts (across the selected marker genes) less than 1,000 were considered as failed due to low sequencing depth. Bulk RNA-seq beta samples with beta deconvolution percentage < 70% and alpha samples with alpha deconvolution percentage < 85% were flagged as failing the purity checks.

For each cell type, we selected donors with bulk RNA-seq data passing the purity checks and for being heterozygous at the exonic (*G6PC2*) SNP rs2232328 according to whole genome sequence analyses. There were seven (non-diabetic) such donors for alpha cells (HPAP004, HPAP017, HPAP052, HPAP075, HPAP083, HPAP090, HPAP097). Reads were aligned to hg38 with STAR v2.7.9a(*64*). Alignments overlapping rs2232328 were extracted and used as input to the GATK v4.2.4.1 ASEReadCounter, which was run with the option –DF NotDuplicateReadFilter. Only two donors with at least 90 reads covering this SNP and of European ancestry were retained (HPAP017 and HPAP075). The null hypothesis of alternate allele frequency=0.5 was tested via a binomial test.

## Acknowledgements

This manuscript is dedicated to the memory of Dr. Franz Matschinsky, a giant in the field and our long-time mentor. We thank Dr. Doris Stoffers for her helpful advice on the project and for suggesting the Phloridzin experiment. We acknowledge and appreciate the efforts of nPOD (https://www.jdrfnpod.org/) for its role in obtaining donor organs utilized in these studies. We thank the Functional Genomics Core of the Penn Diabetes Research Center (P30-DK19525) for next generation sequencing. This work was supported by NIH grants U01-DK134995, UM-1-DK126194, UC4-DK112217, U01-DK-123594, UC4-DK-112232, and U01-DK-123716, the Ruth L. Kirschstein National Research Service Award (NRSA) Individual Predoctoral Fellowship (F31 DK126231-01) to VB, and the U.S-Israel Binational Science Foundation (2019314). BG and DA are supported by grants from the Israel Science Foundation and JDRF (1782/18, 2982/20) and the U.S.-Israel Binational Science Foundation (2019314).

## Supplementary Figures and Tables

**Supplementary Fig. S1:**
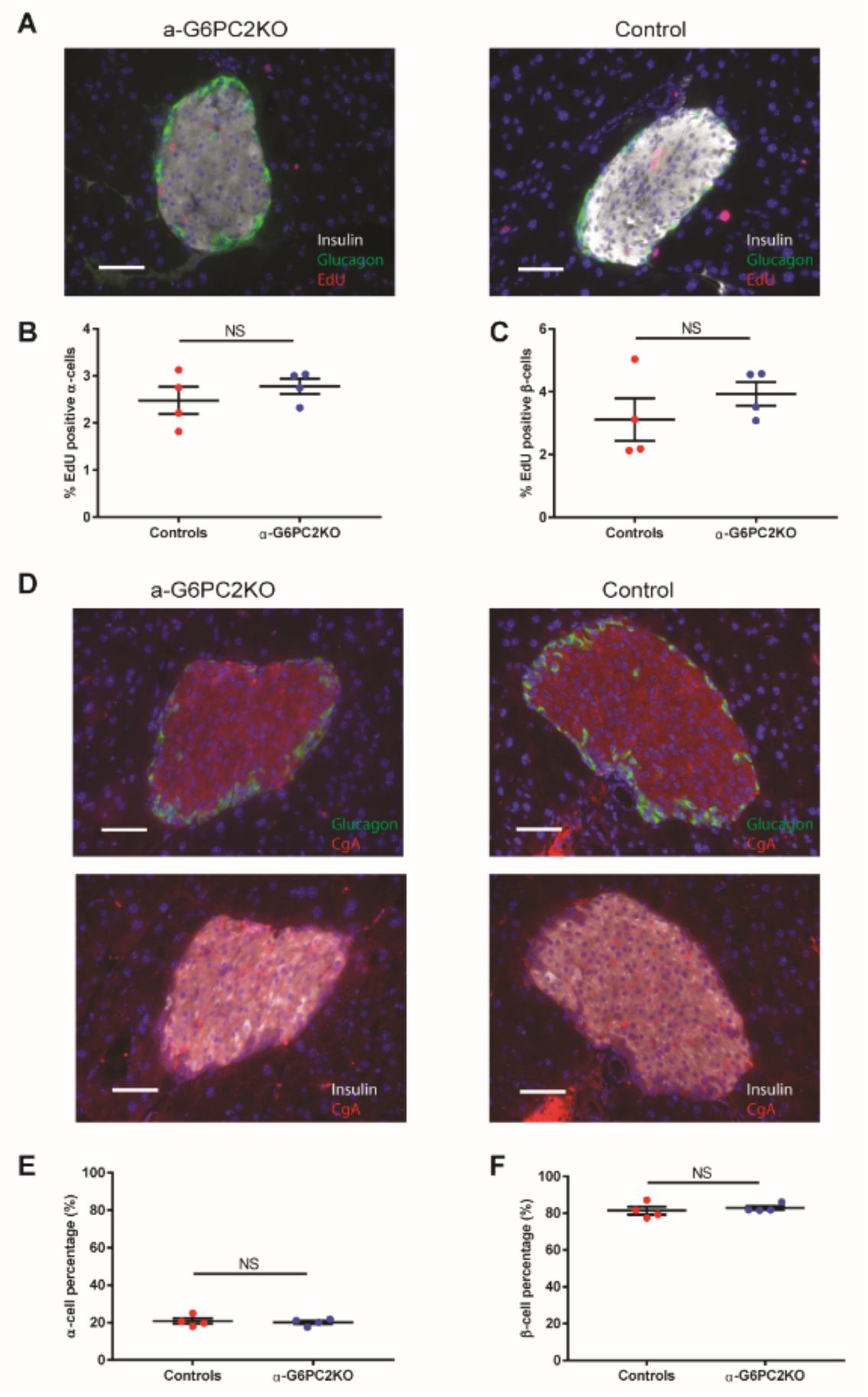
12-week-old α-G6PC2KO mice have normal islet morphology. (**A)** Insulin, glucagon, and EdU shown by co-immunofluorescence (40X). Scale bar: 50μm. **(B)** Glucagon^+^, EdU^+^ cells as percentage of total Glucagon^+^ cells (n=4 for each genotype). **(C)** Insulin^+^, EdU^+^ cells as percentage of total Insulin^+^ cells (n=4 for each genotype). **(D)** Insulin, Glucagon, and CgA shown by co-immunofluorescence (40X). Scale bar: 50μm. **(E)** Glucagon^+^, CgA^+^ cells as percentage of total CgA^+^ cells (n=4 for each genotype). **(F)** Insulin^+^, CgA^+^ cells as a percentage of total CgA^+^ cells (n=4 for each genotype).

**Supplementary Fig. S2:**
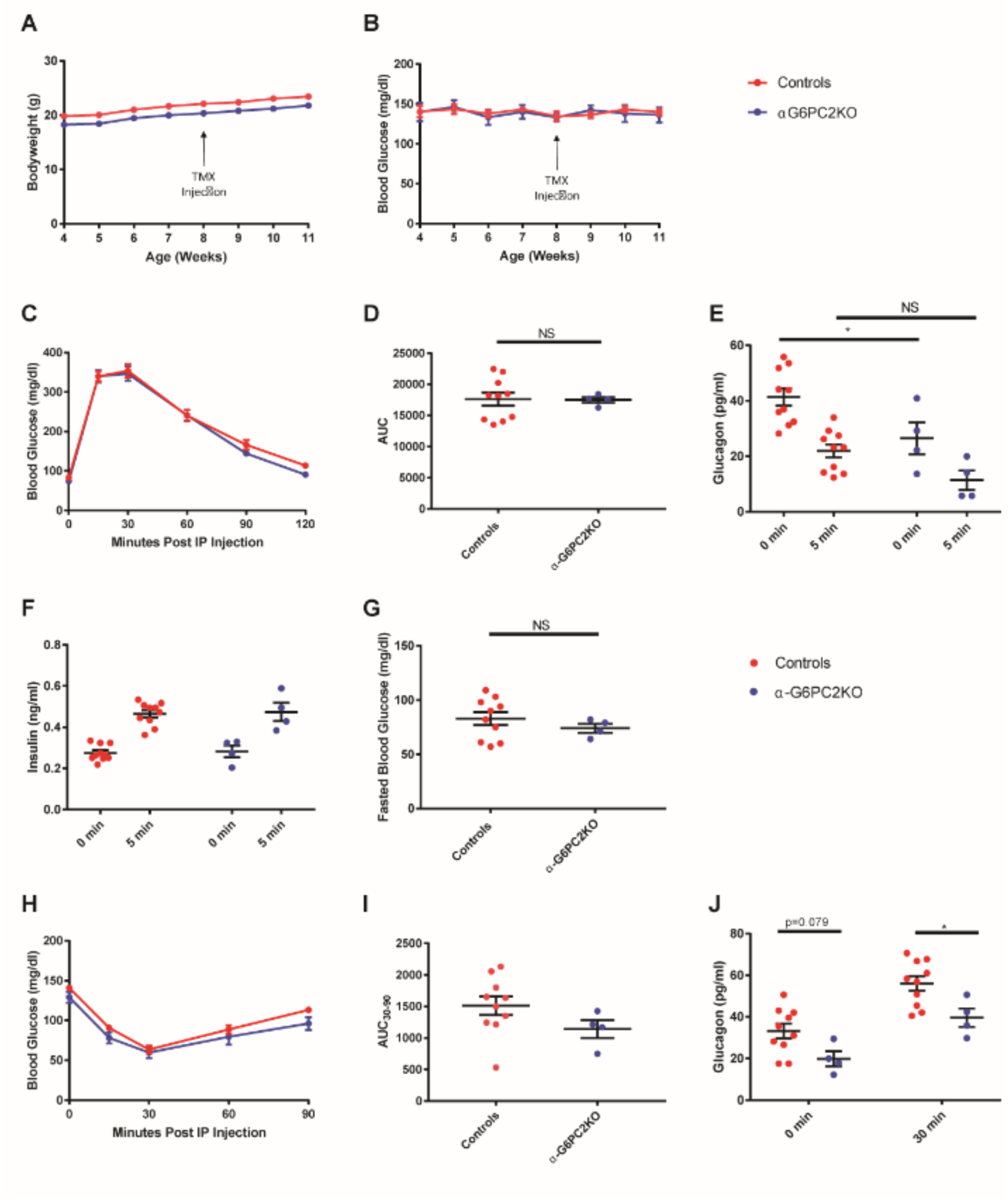
α-cell G6PC2 ablation alters glucose homeostasis in adult female mice. (**A)** Body weights of female α-G6PC2KO and control mice (n=4 for α-G6PC2KO, n=10 total for *G6pc2^loxP/loxP^* and *Gcg-CreER^T2^)*. **(B)** *Ad libitum* blood glucose for female α-G6PC2KO and control mice (n=4 for α-G6PC2KO, n=10 total for *G6pc2^loxP/loxP^* and *Gcg-CreER^T2^)*. **(C)** Intraperitoneal glucose tolerance test (1mg/g bodyweight) (n=4 for α-G6PC2KO, n=10 total for *G6pc2^loxP/loxP^* and *Gcg-CreER^T2^)*. **(D)** Area Under the Curve (AUC) from intraperitoneal glucose tolerance test in (C). **(E)** Plasma glucagon and **(F)** plasma insulin in mice fasted or 5 min after glucose injection in (C). **(G)** Fasted blood glucose measurements from intraperitoneal glucose tolerance test in (C). **(H)** Intraperitoneal insulin tolerance test (0.75U/kg bodyweight) (n=4 for αG6PC2KO, n=10 total for *G6pc2^loxP/loxP^* and *Gcg-CreER^T2^)*. **(I)** Area Under the Curve between 30 and 90 min post-insulin injection (AUC_30-90_) from intraperitoneal insulin tolerance test in (H). **(J)** Plasma glucagon in mice fasted or 30 min after insulin injection in (H). P-values from unpaired Student’s t-test or two-way ANOVA with *post hoc* Bonferroni test: *p < 0.05.

**Supplementary Fig. S3:**
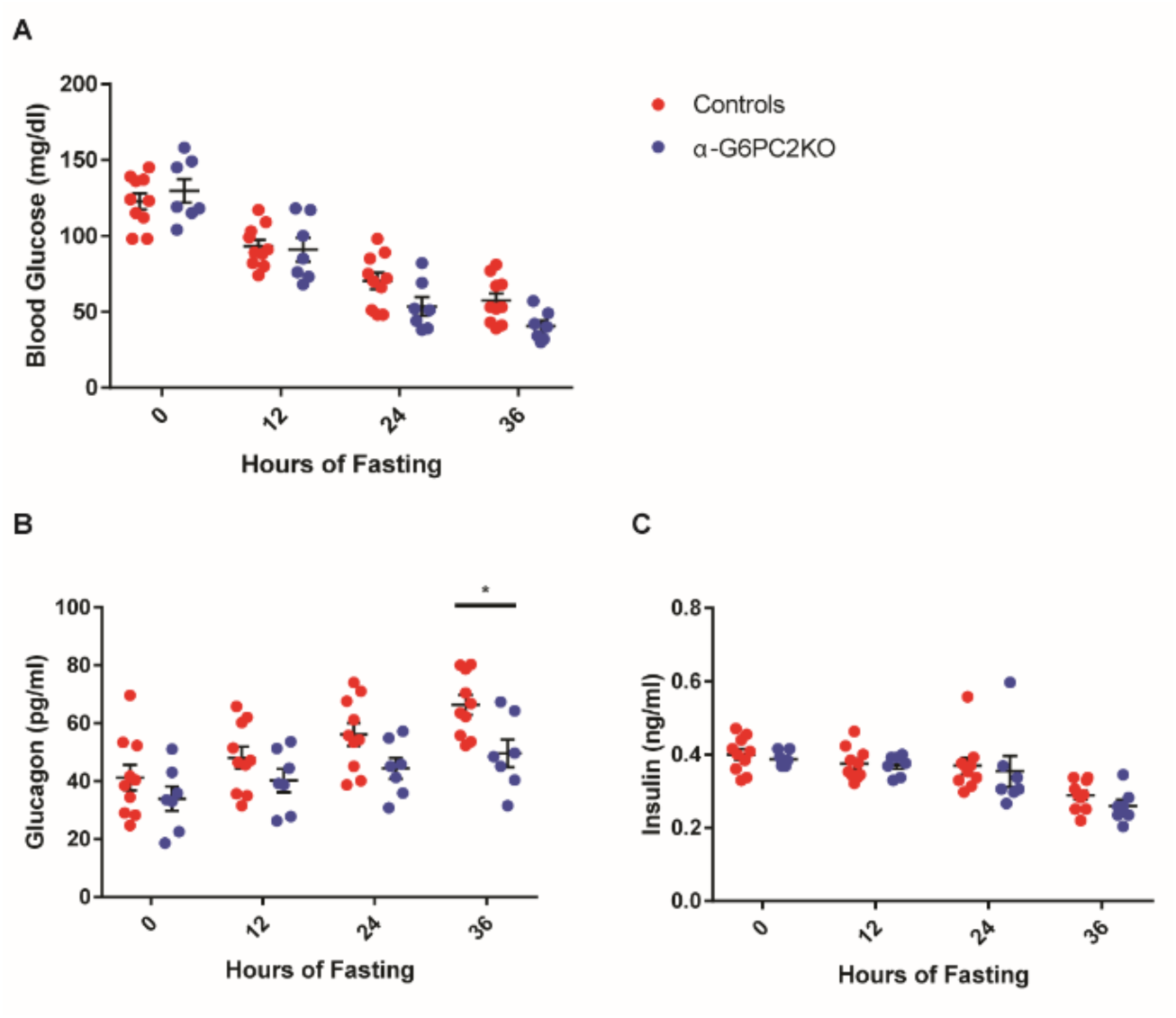
Female α-G6PC2KO mice present lower glucagon levels in response to prolonged fasting-induced hypoglycemia. (**A)** Blood glucose, **(B)** Plasma Glucagon, and **(C)** Plasma insulin for female α-G6PC2KO and control mice (n=7 for α-G6PC2KO, n=10 total for *G6pc2^loxP/loxP^* and *Gcg-CreER^T2^*) during the 36-hour prolonged fast. P-values from two-way ANOVA with *post hoc* Bonferroni test: *p < 0.05.

**Supplementary Fig. S4:**
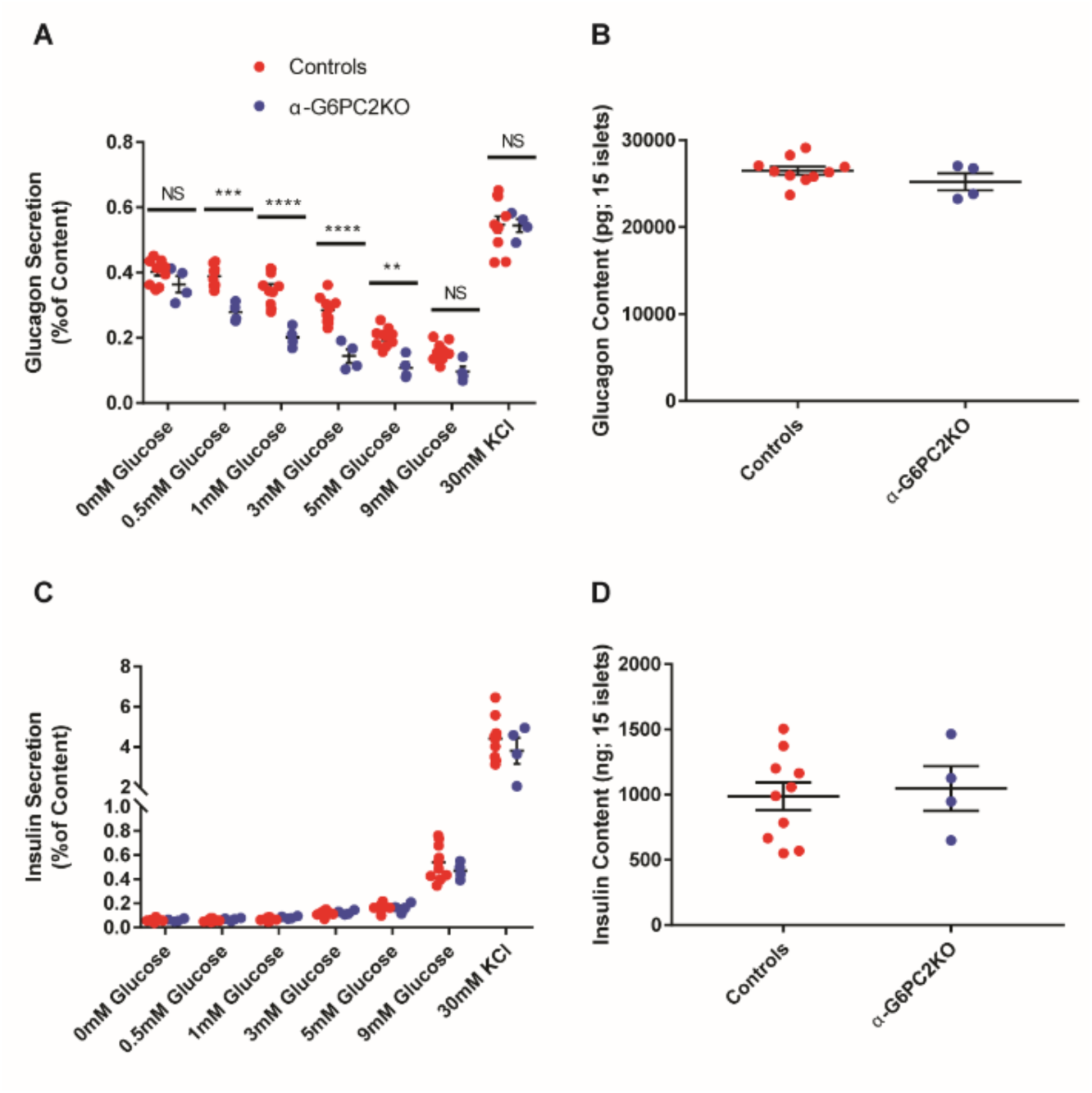
Glucose is more potent in suppressing glucagon secretion in isolated pancreatic islets from female α-G6PC2KO mice. (**A)** Glucagon secretion and **(B)** content from isolated pancreatic islets in the presence of the indicated glucose concentrations with 4mM amino acid mixture or 30mM KCl. **(C)** Insulin secretion and **(D)** content from isolated pancreatic islets in the presence of the indicated glucose concentrations with 4mM amino acid mixture or 30mM KCl. P-values from two-way ANOVA with *post hoc* Bonferroni test: **p<0.01, ***p < 0.001, ****p < 0.0001.

**Supplementary Table S1.**
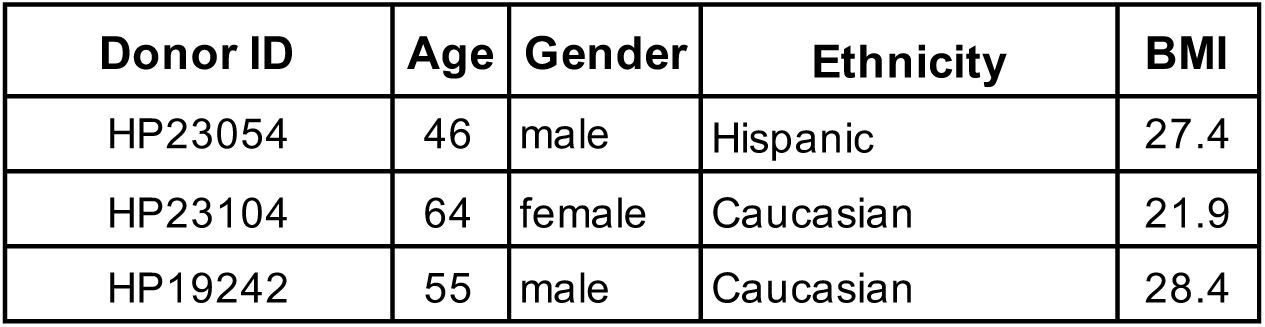
donor information.

| Donor ID | Age | Gender | Ethnicity | BMI |
| --- | --- | --- | --- | --- |
| HP23054 | 46 | male | Hispanic | 27.4 |
| HP23104 | 64 | female | Caucasian | 21.9 |
| HP19242 | 55 | male | Caucasian | 28.4 |

